# Elucidation of small molecule passive permeation across lipid membranes using conventional solution state NMR methods

**DOI:** 10.1101/2022.10.01.510446

**Authors:** Angela Serrano-Sanchez, Joseph Cassar, Lisa J. White, Precious I. A. Popoola, Jennifer R. Hiscock, Jose L. Ortega-Roldan

**Affiliations:** School of Biosciences. University of Kent, Canterbury, UK. CT2 7NJ; School of Chemistry and Forensic Science. University of Kent, Canterbury, UK. CT2 7NH

**Keywords:** membrane permeability, antimicrobials, Solution NMR, membrane interaction

## Abstract

Quantifying small molecule uptake across a biological membrane in any cell system is crucial for the development of efficacious and selective drugs. However, obtaining such data is not trivial, especially in bacterial systems. Herein, we present an assay which enables the determination of the degree of passive permeation and membrane interaction of mixtures of small molecules in vesicles of a desired lipid composition, including that of bacterial membranes. The assay employs highly accessible conventional solution NMR experiments, exploiting the paramagnetic relaxation enhancement effect, and allows the measurement of membrane permeation on mixtures of any number of small molecules which do not exhibit heterogeneous molecular signal overlap in under 20 minutes. As a proof-of -principle we apply this methodology to candidates from a class of supramolecular self-associating amphiphiles, members from which have been shown to interact with biological phospholipid membranes and elicit an antimicrobial effect, allowing the determination and comparison of their membrane permeability and membrane interaction properties.

## Introduction

Antimicrobial resistance (AMR) is one of the greatest global health threats, with a recent study reporting that in 2019 a greater number of people died from the primary effects of AMR than malaria or HIV/AIDS^[1]^. Therefore, novel therapeutic agents with innovative modes of action are required to address this major health threat. We believe that a significant limitation for novel antimicrobial agents translating into the clinic maybe related to the inability of these compounds to reach a site of action within the organism itself due to penetrate to the organism and achieve their site of action^[2]^. One of the biggest challenges in developing antibiotics against multi-drug resistant bacterial pathogens is their small-molecule uptake as their membrane composition differ substantially from mammalian cells membranes^[3,4]^, and therefore, assays able to monitor small molecule uptake in any lipid composition are required.

Currently, several in vitro experimental models provide a permeability evaluation and transport of pharmaceutical molecules across at least one cellular membrane to reach their target^[5]^. Among the high throughputs in vitro techniques, those based in artificial membranes such as parallel artificial membrane permeability assay (PAMPA)^[6]^, and the cell based Caco-2 assay^[7]^ are the most widely used. These models are important in the initial stages of drug discovery in mammalian cells, but are not available for bacterial systems, do not allow therapeutic mode of action or selectivity analysis, and typically involve long assay times, cell culturing and specialised equipment, hence resulting in an elevated cost.

Herein, we present an accessible, novel approach able to assess small molecule passive permeability through vesicles composed of any type of lipid composition using solution NMR spectroscopy, widely accessible in chemistry and biological laboratories. This methodology is considerably quicker (1 mixture of small molecules/20 minute) and allows the study of any membrane composition in an affordable way.

We have used this methodology to compare the permeabilities and degree of membrane interaction of a set of reference compounds, glucose, which cannot permeate through the membrane, and the membrane permeable indole, and a set of test compounds from the Supramolecular Self-associating Amphiphile (SSA) family ^[8–10]^(Figure 1). This technology has already shown antimicrobial efficacy against clinically relevant Gram-positive methicillin resistant *Staphylococcus aureus* (MRSA) and Gram-negative *Escherichia Coli* (*E. coli*), it exhibits a druggable profile in vivo, and enhances antimicrobial and anticancer activities of different commercial drugs^[11,12]^.

**Figure 1:**
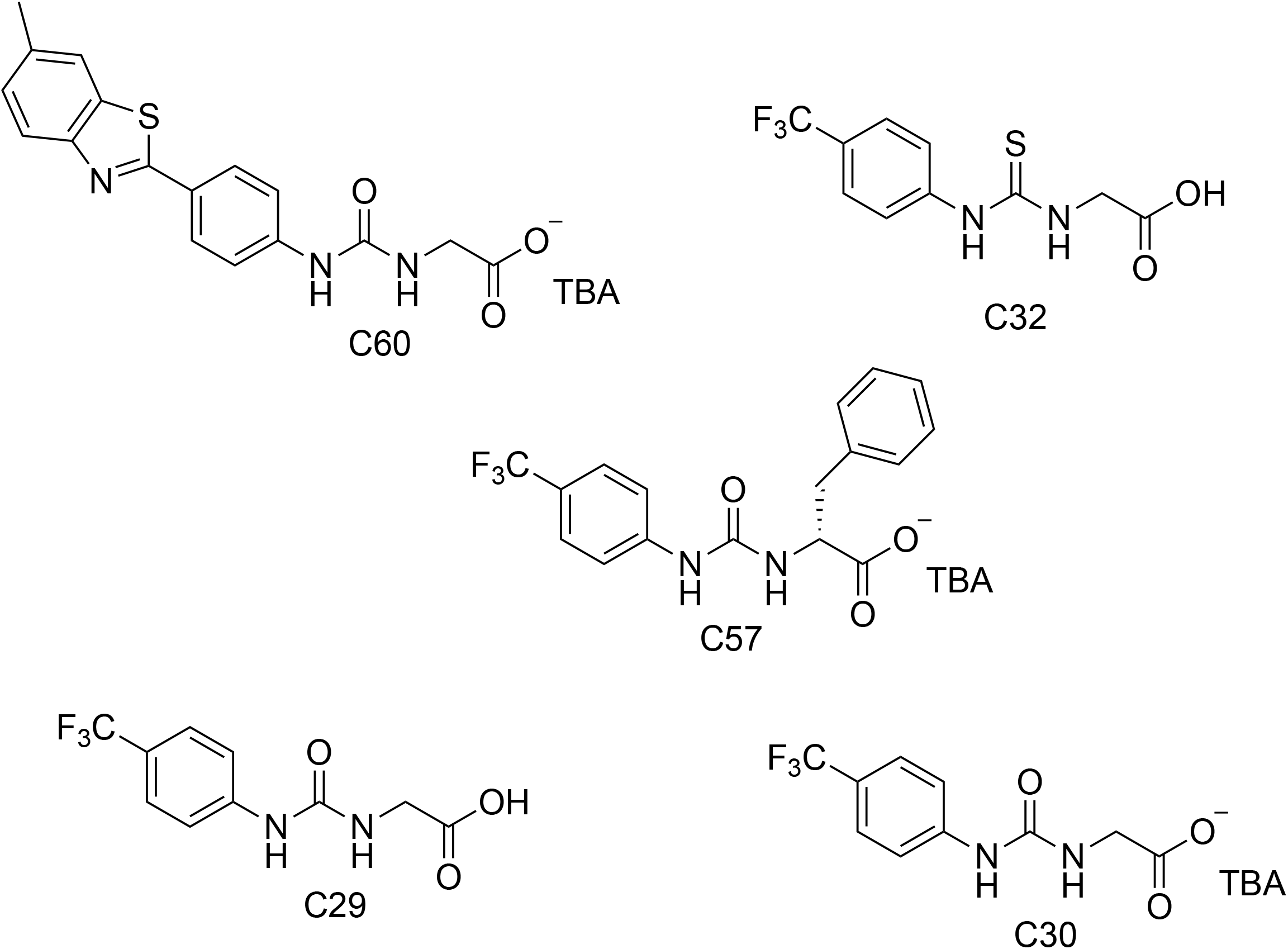
Chemical structures of SSAs C29, C30, C32, C57 and C60. TBA = Tetrabutylammonium.

Proton NMR T1 and T2 relaxation rates depend, amongst other parameters, on the rotational correlation time of the molecule. For larger molecules which are characterized by longer T1 values this that can modulate the intensity of ^1^H NMR signals if the sum of the acquisition time and the relaxation delay is not long enough to return to equilibrium during the relaxation delay in a multi transient experiment. The addition of solvent PRE reagents has been shown to induce a much larger reduction of T1 than T2 values for small molecules^[13]^. However, the measured effect of the PRE on T2, which lead to a decrease in the ^1^H intensities, can be maximized employing CPMG pulse sequences^[14]^.

We sought to test if the addition of a solvent PRE reagent only in the outside compartment of a vesicle will exert a different effect in the ^1^H NMR peak intensities of molecules with different permeation rates due to the different modulation of T1 and T2 values. We measured ^1^H 1D CPMG spectra for the non membrane-permeable glucose and the highly membrane-permeable indole in the absence and presence of vesicles prepared with the gram negative model bacteria *E. coli* lipids with and without the PRE reagent Mn^2+^. Three different CPMG spin-lock times, 20, 50 and 150 mS, were used to maximise the PRE effect on T2, and hence improve differentiation between molecules with different permeation rates.

Glucose, which cannot permeate through the membrane unaided, shows a significant drop of intensity of its ^1^H resonances in the presence of Mn^2+^ both in the presence and absence of lipid vesicles, consistent with PRE-induced broadening of the signals. Indole, on the other hand, showed an opposite effect. While in the absence of vesicles Mn^2+^ induced line broadening, in the presence of vesicles we saw the opposite effect, with an increase in the signal intensities upon addition of Mn^2+^. As previously discussed, this effect can be attributed to the reduction of T1 values of soluble indole molecules which have spent some time associated with the lipid vesicle (Figure 2) counteracting the effects of the increase in T1 due to the molecular mass of the vesicle-indole complex during the CPMG period.

**Figure 2:**
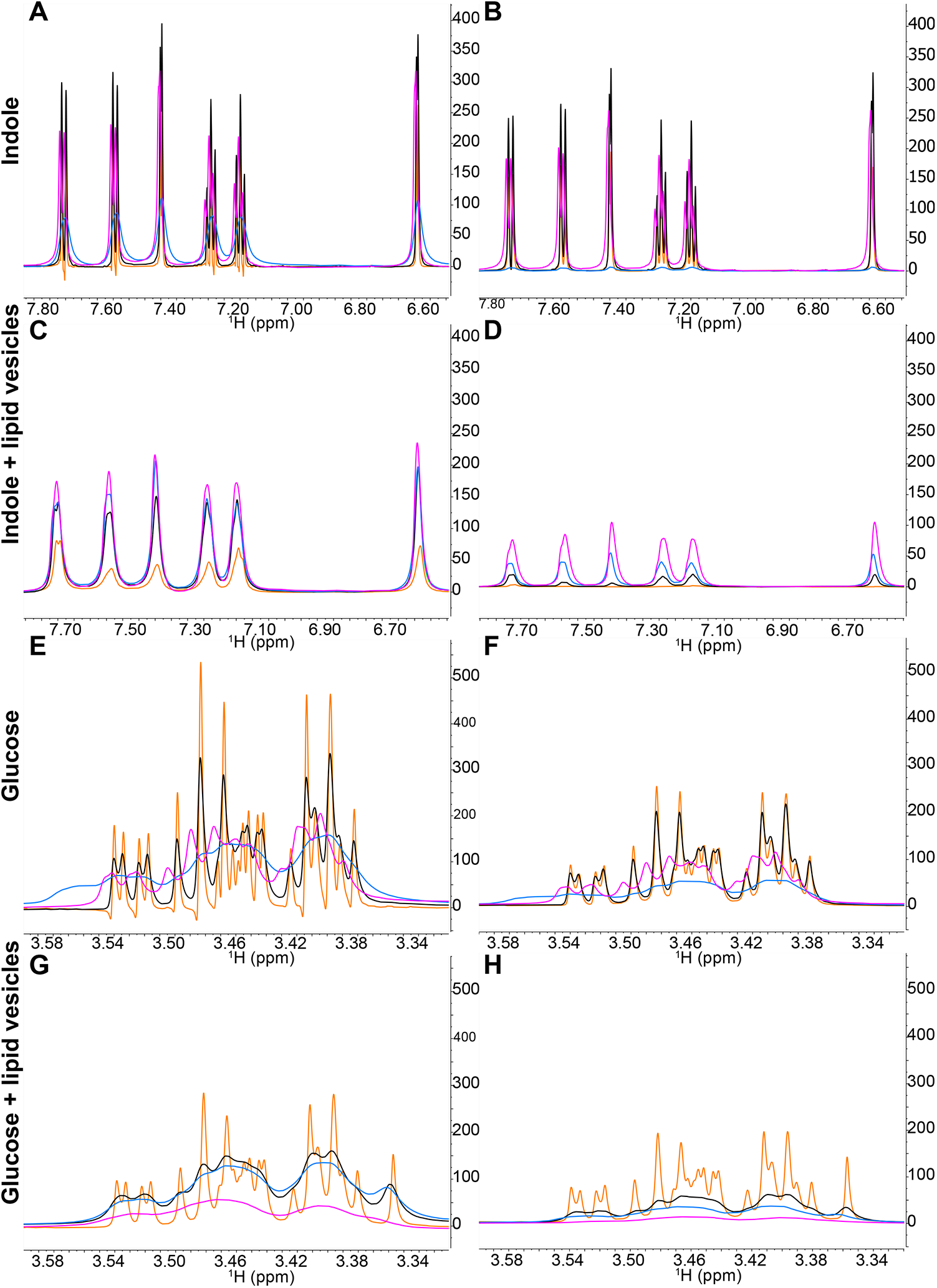
Proton 1D CPMG NMR spectra collected with a spin-lock time of 20 mS (left column (A, C, E and G) and 150 mS (right column (B, D, F and H) of the reference compounds indole and glucose in the absence and presence of *E. coli* lipid vesicles at increasing concentrtations of Mn^2+^ (0 mM (orange); 0.5 mM (black); 2 mM (blue); 4 mM (magenta)). Samples contained 4mM glucose and 2mM indole ± 1.1 mg/mL vesicles ± increasing concentrations of MnCl_2._

The effect of permeation rates on the PRE modulation of 1H intensities can be parametrised by comparing the intensity ratios of ^1^H resonances in ^1^H 1D NMR CPMG experiments with the same spin lock times with and without Mn^2+^ in the presence and absence of lipid vesicles. A value of 1 implies no detectable permeation through the vesicles and values higher than one imply detectable permeation through the vesicles (Equation 1).

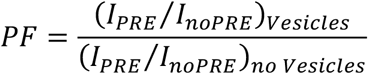

**Equation 1**. The Permeability Factor (PF) depends on the intensity ratio of ^1^H 1D CPMG NMR signals of small molecules in the presence and absence of vesicles and a solvent PRE reagent.

This permeability factor, and its change with different CPMG times, can be used to compare the permeability of different compounds in vesicles of the same composition or to assess the permeability of a single compound in vesicles with different lipid compositions (i.e. extracted from different cell strains) (Table 1).

**Table 1.** Permeability and membrane interaction factors of Glucose, Indole and a subset of compounds from the SSA family. PF and MIF values were calculates using Equations 1 and 2, respectively. All values given with a ±0.1 uncertainty unless specified in the table.

| Compound | PF<br>(20ms) | PF<br>(50ms) | PF<br>(150ms) | MIF<br>(20ms) | MIF<br>(50ms) | MIF<br>(150ms) |
| --- | --- | --- | --- | --- | --- | --- |
| Glucose | 0.8 | 0.5 | 0.4 | 0.8 | 0.8 | 0.8 |
| Indole | 2.3 | 4.5 | 8.6 | 0.3 | 0.1 | 0.0 |
| C29 | 0.8 | 0.7 | 0.5 | 1.0 | 1.0 | 0.9 |
| C30 | 1.1 | 1.2 | 1.1 | 1.1 | 1.1 | 1.0 |
| C32 | 1.4 | 1.2 | 0.9 | 1.0 | 1.0 | 0.9 |
| C57 | 0.9 | 1.0 | 1.0 | 0.9 | 0.9 | 0.9 |
| C60 | 5.3±0.2 | 5.5±0.3 | 6.6±1.2 | 0.6 | 0.4 | 0.2 |

Similarly, the intensity ratio between ^1^H resonances in ^1^H 1D NMR CPMG experiments with the same spin lock times in the presence and absence of lipid vesicles reports on the degree of interaction between small molecules and the membrane. A value of 1 signifies no detectable membrane interaction and a value below 1 indicates detectable membrane interaction (Equation 2), with the affinity being related to the CPMG spin-lock time at which the deviation from 1 is observed (Table 1).

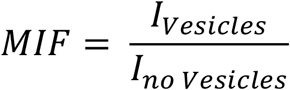

**Equation 2**. The Membrane Interaction Factor (MIF) depends on the intensity ratio of ^1^H 1D CPMG NMR signals of small molecules in the presence and absence of vesicles.

It is important to note that PF and MIF values rely on intensities extracted from unbound ligand in the ^1^H 1D NMR CPMG spectra, and therefore the molecules studied must not fully self-associate in solution at the concentrations used for the assay. The degree of molecular self-association can be obtained easily through comparing the intensity ratio between ^1^H 1D NMR CPMG spectra collected at 150mS and 20mS spin-lock times, with a value of 1 indicating no self-association and a value below 1 indicating the presence of self-association.

We subsequently used this methodology to assess and classify the permeation rates and membrane-interaction affinities of the control substances indole and glucose and a set of 5 representative compounds from a novel class of supramolecular self-associating amphiphile (SSAs), using vesicles prepared with the same *E. coli* lipids as those prepared previously for the control experiments (Figure 3). The hypothesised basis for the therapeutic activity of this class of related compounds includes their ability to selectively coordinate with phospholipids of different head group composition and permeate into the cell^[15,16]^.

**Figure 3:**
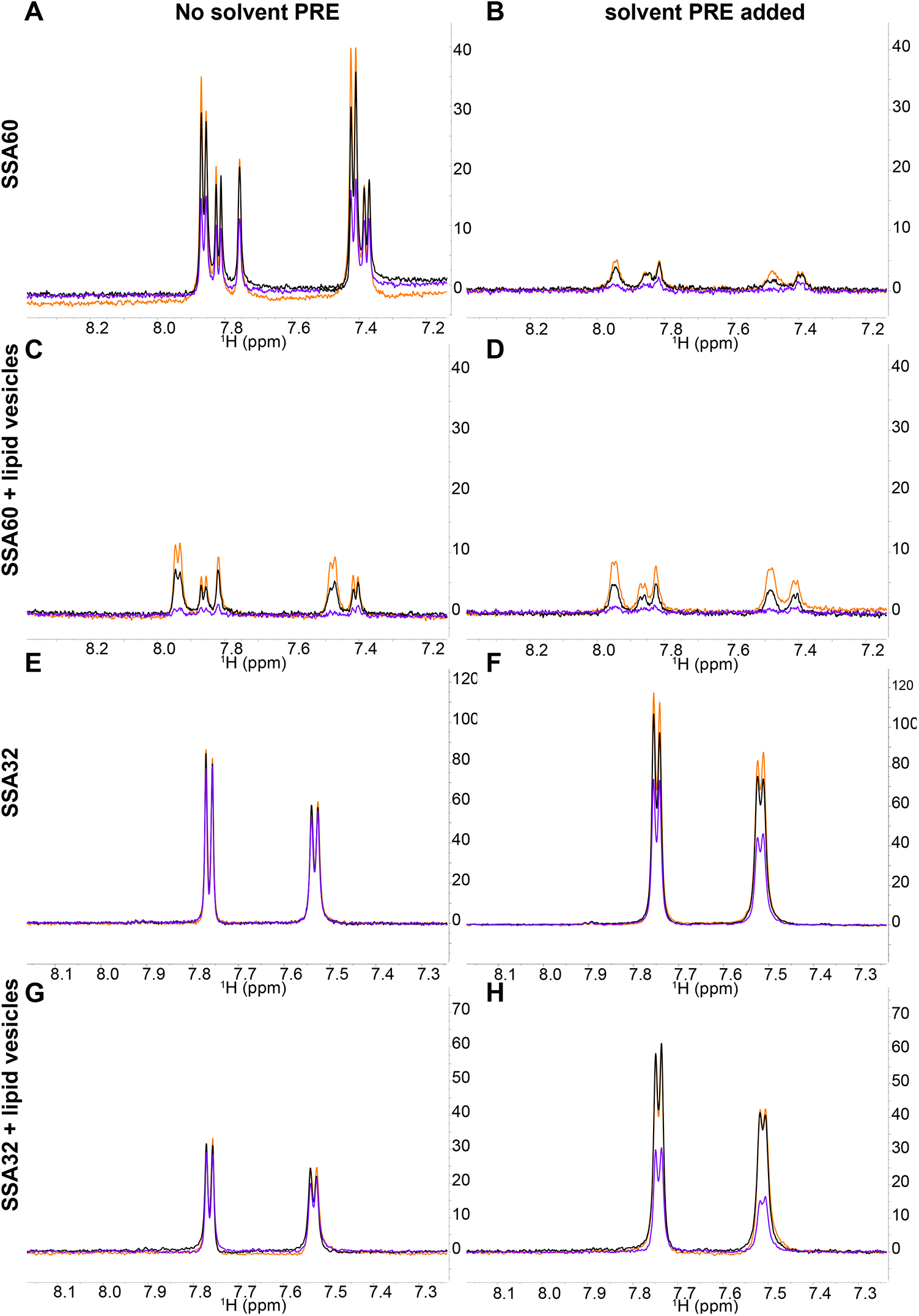
Proton 1D CPMG NMR spectra collected with a spin-lock time of 20, 50 and 150 mS (orange, black and purple, respectively) of SSA60 (A-D) and SSA32 (E-H) in the absence (A-B, E-F) and presence (C-D, G-H) of *E. coli* lipid vesicles without (A, C, E, G) and with (B, D, F, H) the addition of 0.5M Mn^2+^ (left and right column, respectively).Samples contained 200uM of the SSA compounds ± 1.1 mg/mL vesicles ± 0.5 mM MnCl_2_

The intensity ratios of all aromatic peaks of indole and SSA compounds and the aliphatic peaks of glucose, collected in the absence or presence of lipid vesicles and Mn^2+^, were used for the permeability analysis. The permeability and membrane interaction factors for these molecules were calculated using Equations 1 and 2 (Table 1). As expected, indole showed high permeability factors (PF) and low membrane interaction factors (MIF), indicative of high permeability and high membrane interaction affinity. Glucose, on the other hand, showed permeability factors below 1, consistent with its lack of membrane permeability, and MIF of 0.8, indicating a weak membrane interaction affinity consistent with previous findings^[17]^. Significant differences can be observed in the permeability of SSA compounds. C60 showes the highest PF and MIF, resembling the properties of indole. C32 shows some slight permeability, although no detectable membrane interaction, an indication of its low affinity for *E. coli* lipid vesicles. C57 shows no detectable permeation, but noticeable membrane interaction. C29 and C30 show no detectable membrane permeation or interaction.

## Discussion

Herein, we present a new assay using solution NMR, a methodology accessible to most chemistry and biological laboratories, able to determine the membrane permeability and degree of membrane interaction of small molecules. The method exploits the NMR solvent PRE effect using CPMG experiments, and the ability of membrane permeable molecules to travel to the interior of lipid vesicles, shielding themselves from the solvent PRE reagent. The assay consists in the measurement of ^1^H 1D CPMG NMR spectra in four different samples all containing the mix of drugs: 1 – without vesicles, 2 – without vesicles with 0.5mM Mn2^+^ added to the buffer, 3 – with vesicles, and 4 – with vesicles with 0.5mM Mn2^+^ added to the buffer.

This method relies on the elevation of ^1^H 1D CPMG intensities for molecules permeating through the membrane, which is caused by the bidirectional equilibrium into and out of the vesicles due to the lack of asymmetry in the lipid composition of the vesicles used. As a result, these molecules spend a large time bound to the vesicle, hence the increase in intensity due to PRE effects on T1, and they are partially protected from the PRE effect on T2 in the extravesicular space. A similar effect would result if a molecule was only able to interact with the membrane but could not permeate, likely due to interactions with the headgroup region of the lipids. This region is in close contact with the solvent, and therefore is more exposed to the solvent PRE. To assess if the molecules under study are permeating or only interacting with the membrane, a comparison between the MIF, which is obtained in the absence of PRE, and the PF is needed. C60 shows a high PF and MIF values in the range of 0.6-0.2, while indole shows equally high PF and MIF values of 0.3 – 0. The higher MIF values of C60 suggest that these compounds are able to permeate more that indole, and that indole has a higher affinity to the lipid membrane and stays closely associated to it.

Glucose, a membrane-impermeable molecule was also used as a control. While indole showed high PF and low MIF values, glucose showed low PF and values of MIF close to 1. The deviations found for glucose from the ideal values for a non-permeable molecule (PF and MIF =1) arise from a degree of self-association, as indicated by the intensity ratio between glucose peaks in CPMG experiments collected with low (20mS) and high (150mS) spinlock times in the absence of vesicles and solvent PRE. Self-association needs to be measured and considered for accurate PF measurements, as high degrees of self-association would lower PFs and hinder the detection of permeation.

SSAs are amphiphilic compounds and thus could present surfactant properties. To explore the interaction of SSAs with all types of lipid membranes, Boles and co-workers explored the relation between membrane lysis/permeation and interaction values to support the hypothesis that the increase in antimicrobial efficacy depends on the general surfactant properties of the SSA compound^[15]^. Herein, we can also analyse the relation between permeation and membrane interaction with the lysis of lipid vesicles obtained by the previous study. Their results revealed that C48 could not disrupt the lipid membrane from *E. coli* which supports our finding that this compound did not permeate the lipid vesicles in our assay. C30 presented slight permeability while C60 presented very high lipid permeability in *E. coli* lipid vesicles. Both C30 and C60 presented lysis properties against all the phospholipid membranes, markedly higher in C60, showing a good correlation between the permeability values obtained in our NMR-based assay and the efficacy of these antimicrobials due to their ability to disrupt cell membranes.

In conclusion, we present a high throughput, accessible and low-cost assay to assess the permeation and degree of membrane interaction of small molecules. The assay can be used with any lipid composition, including lipids extracted from natural sources, allowing permeation studies of antimicrobials, and can assess the permeability of any mixture of small molecules with non-overlapping ^1^H NMR peaks in as little as 20 minutes.

## Acknowledgements

J. Hiscock would also like to thank UKRI for the funding of her Future Leaders Fellowship (MR/T020415/1). J. Ortega-Roldan would like to thank Dr Gary Thompson (University of Kent) and Dr Lorena Varela for NMR training, useful discussion, and/or manuscript revision.

